# Heterogeneity of morphometric similarity networks in health and schizophrenia

**DOI:** 10.1101/2024.03.26.586768

**Authors:** Joost Janssen, Ana Guil Gallego, Covadonga M. Díaz-Caneja, Noemi González Lois, Niels Janssen, Javier González-Peñas, Pedro M. Gordaliza, Elizabeth E.L. Buimer, Neeltje E.M. van Haren, Celso Arango, René S. Kahn, Hilleke E. Hulshoff Pol, Hugo G. Schnack

## Abstract

**Introduction:** Morphometric similarity is a recently developed neuroimaging phenotype of inter-regional connectivity by quantifying the similarity of a region to other regions based on multiple MRI parameters. Altered average morphometric similarity has been reported in psychotic disorders at the group level, with considerable heterogeneity across individuals. We used normative modeling to address cross-sectional and longitudinal inter-individual heterogeneity of morphometric similarity in health and schizophrenia.

**Methods:** Morphometric similarity for 62 cortical regions was obtained from baseline and follow-up T1-weighted scans of healthy individuals and patients with chronic schizophrenia. Cortical regions were classified into seven predefined brain functional networks. Using Bayesian Linear Regression and taking into account age, sex, image quality and scanner, we trained and validated normative models in healthy controls from eleven datasets (n = 4310). Individual deviations from the norm (z-scores) in morphometric similarity were computed for each participant for each network and region at both timepoints. A z-score ≧ than 1.96 was considered supra-normal and a z-score ≦ -1.96 infra-normal. As a longitudinal metric, we calculated the change over time of the total number of infra- or supra-normal regions per participant.

**Results:** At baseline, patients with schizophrenia had decreased morphometric similarity of the default mode network and increased morphometric similarity of the somatomotor network when compared with healthy controls. The percentage of patients with infra- or supra-normal values for any region at baseline and follow-up was low (<6%) and did not differ from healthy controls. Mean intra-group changes over time in the total number of infra- or supra-normal regions were small in schizophrenia and healthy control groups (<1) and there were no significant between-group differences.

**Conclusions:** In a case-control setting, a decrease of morphometric similarity within the default mode network may be a robust finding implicated in schizophrenia. However, normative modeling suggests that significant reductions and changes over time of regional morphometric similarity are evident only in a minority of patients.

## Introduction

Schizophrenia is a severe psychiatric disorder that affects about 1% of the global population (Jauhar et al., 2022). Despite decades of research, the prognosis in a substantial proportion of patients remains poor (Jauhar et al., 2022). A considerable challenge hindering the development of more effective, personalized treatment strategies for schizophrenia lies in the unresolved neurobiological heterogeneity that characterizes the disease (Alnæs et al., 2019; Brugger and Howes, 2017). While morphological reductions in brain volume and cortical thickness may be relatively robust at the group level in schizophrenia, there is considerable variation across individuals (Brugger and Howes, 2017; Haijma et al., 2013; Kelly et al., 2018; Segal et al., 2023; van Erp et al., 2018). The normative modeling framework aims to address the issue of individual phenotypic heterogeneity by establishing normative standards for neurobiological variables and subsequently assessing individual’s deviation from these norms (Marquand et al., 2019). Studies applying normative model analysis in schizophrenia are relatively sparse but have shown that abnormal deviations of regional cortical thickness and volume are frequent at the individual level but rarely occur consistently in the same locations and with the same severity across individuals (Di Biase et al., 2022; Lv et al., 2021; Segal et al., 2023; Wolfers et al., 2018).

Morphometric similarity is a recently developed neuroimaging phenotype of cortical inter-regional connectivity by quantifying the similarity of a region to all other regions based on multiple MRI parameters assessed at each region (Sebenius et al., 2023; Seidlitz et al., 2018). That is, each brain region is represented as a vector of several MRI features, such as cortical thickness and volume, and based on the pairwise correlation between the regional feature vectors, morphometric similarity can be estimated.

Morphometric similarity has been validated as a proxy for inter-regional connectivity, as regions with high morphometric similarity show stronger axonal connectivity (Seidlitz et al., 2018). In contrast to cortical volume and thickness, little is known about the typical age-dependency of morphometric similarity between late childhood and old age. Normative modeling in a large sample of healthy controls with a wide age range provides an excellent opportunity to characterize regional differences in the age-dependency of typical morphometric similarity (Zhukovsky et al., 2022). In psychosis, morphometric similarity has shown recent promise as a relevant phenotype because findings of reduced average morphometric similarity in frontal and temporal regions have been replicated in three independent groups of adult individuals with psychosis and one group of individuals with early-onset schizophrenia (Morgan et al., 2019; Yao et al., 2023). In addition, a disproportionate number of regions that showed group-average reductions in morphometric similarity in individuals with psychosis were part of the default mode network (DMN) which concurs with DMN deficits reported from functional MRI studies (Broyd et al., 2009; Heuvel and Sporns, 2019; Morgan et al., 2019). However, it is worth noting that diagnostic investigations into morphometric similarity have predominantly relied on cross-sectional group-level averages, thereby overlooking potentially substantial heterogeneity of morphometric similarity at the individual level (Morgan et al., 2019; Yao et al., 2023; Zhukovsky et al., 2022).

Longitudinal assessments of morphometric similarity combined with normative modeling allows not only for cross-sectional quantification of individual deviation but also enables assessment of intra-individual change in deviation over time and its relation to the established normative range (Barbora Rehák Bučková et al., 2024; Di Biase et al., 2023).

Here, we apply a normative modeling framework to morphometric similarity and conduct a longitudinal study of this metric for the first time. In this study, we aim to parse the cross-sectional and longitudinal heterogeneity of morphometric similarity in schizophrenia, by assessing a longitudinal sample of adult individuals with schizophrenia and healthy controls.

## Methods

### Sample

Eleven datasets were combined to create the full sample. These datasets are described in Figure A including the sample size, age, and sex distribution of each dataset. We included healthy participants from ten publicly available datasets: the Amsterdam Open MRI Collection (aomic) (id1000, piop1 and piop2) datasets (Snoek et al., 2021), the Cambridge Centre for Ageing and Neuroscience (camcan) dataset (Shafto et al., 2014), the Dallas Longitudinal Brain Study (dlbs) dataset (Lu et al., 2011), the Information eXtraction from Images (ixi) dataset (http://brain-development.org/ixidataset/), the Narratives (narratives) dataset (Nastase et al., 2021), the Open Access Series of Imaging Studies (oasis3) dataset (Marcus et al., 2010), the NKI-Rockland (rockland) dataset (Nooner et al., 2012) and the Southwest University adult lifespan (sald) dataset (Wei et al., 2018). One dataset is not publicly available and consisted of a large longitudinal sample of individuals with schizophrenia and healthy participants aged 16-68 years (at baseline) from the Utrecht Schizophrenia project and the Genetic Risk and Outcome of Psychosis (GROUP) consortium. From this longitudinal clinical sample we included individuals who had T1-weighted magnetic resonance imaging (MRI) scan acquisitions at baseline and follow-up. Two identical scanners were used and all included participants had their baseline and follow-up scans acquired on the same scanner. Detailed information regarding diagnostic criteria, clinical assessments, MRI acquisition and image quality control assessment of the Utrecht Schizophrenia project and the GROUP consortium are described in (Hulshoff Pol et al., 2001; Janssen et al., 2021; Korver et al., 2012; Kubota et al., 2015). Additional demographic, cognitive, and clinical characteristics of the longitudinal clinical sample are provided in Table A. All participants provided written informed consent. Subject recruitment procedures and informed consent forms, including consent to share de-identified data, were approved by the corresponding institutional review board where data were collected.

**Figure A.**
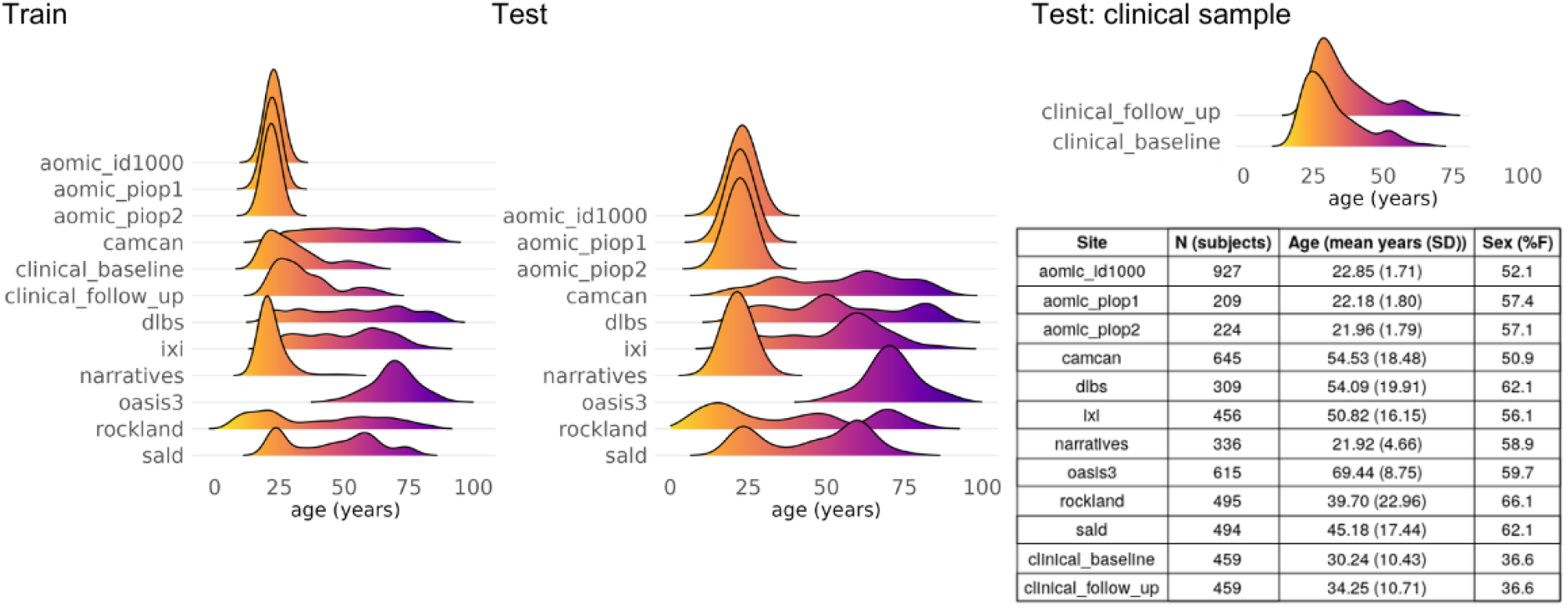
Eleven datasets were used in the study. Ten datasets were cross-sectional and included healthy participants. For each of these ten datasets, 90% of individuals were included in the training set and 10% were part of the test set. One longitudinal clinical dataset (two timepoints) included healthy controls and individuals with chronic schizophrenia. Of the healthy controls belonging to the longitudinal clinical dataset, 20% were included in the training set and 80% in the test set. All individuals with schizophrenia were included in the test set. The age distribution for each dataset within the training/test samples is shown. The table shows the descriptives per dataset. N, number of subjects.

**Table A.**
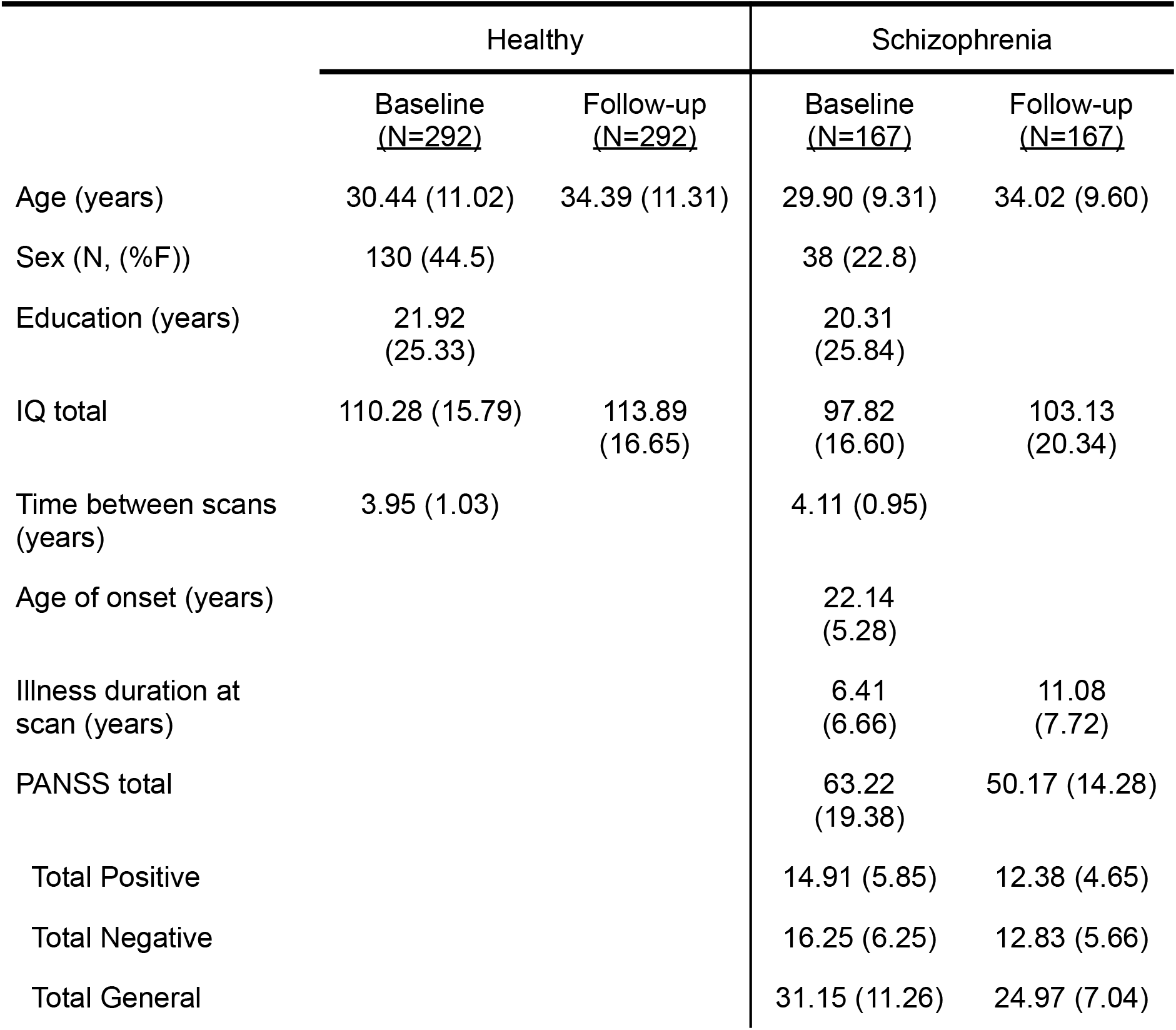
Demographic, cognitive, imaging, and clinical characteristics of the clinical dataset. All descriptors are mean (standard deviation) unless otherwise specified. N, number of subjects; F, female; PANSS, Positive and Negative Syndrome Scale (Kay et al., 1987).

|  | Healthy |  | Schizophrenia |  |
| --- | --- | --- | --- | --- |
|  | Baseline<br>(N=292) | Follow-up<br>(N=292) | Baseline<br>(N=167) | Follow-up<br>(N=167) |
| Age (years) | 30.44 (11.02) | 34.39 (11.31) | 29.90 (9.31) | 34.02 (9.60) |
| Sex (N, (%F)) | 130 (44.5) |  | 38 (22.8) |  |
| Education (years) | 21.92<br>(25.33) |  | 20.31<br>(25.84) |  |
| IQ total | 110.28 (15.79) | 113.89<br>(16.65) | 97.82<br>(16.60) | 103.13<br>(20.34) |
| Time between scans<br>(years) | 3.95 (1.03) |  | 4.11 (0.95) |  |
| Age of onset (years) |  |  | 22.14<br>(5.28) |  |
| Illness duration at<br>scan (years) |  |  | 6.41<br>(6.66) | 11.08<br>(7.72) |
| PANSS total |  |  | 63.22<br>(19.38) | 50.17 (14.28) |
| Total Positive |  |  | 14.91 (5.85) | 12.38 (4.65) |
| Total Negative |  |  | 16.25 (6.25) | 12.83 (5.66) |
| Total General |  |  | 31.15 (11.26) | 24.97 (7.04) |

### Image processing and quality control

All T1-weighted images from all datasets were processed centrally using the FreeSurfer analysis suite (v7.1) with default settings (Fischl, 2012). Estimates of 1) cortical volume, 2) surface area, 3) average cortical thickness, 4) average curvature, and 5) Gaussian curvature were calculated for each of the 62 regions of the Desikan-Killany-Tourneville (DKT) atlas (Klein and Tourville, 2012). The Freesurfer Euler number was extracted as a proxy for image quality (Rosen et al., 2018). Subjects were removed if the maximum, absolute, within-dataset centered Euler number was larger than 10 (Rutherford et al., 2023).

### Morphometric similarity

Using regional measurements of cortical thickness, surface area, cortical volume, mean curvature and Gaussian curvature we calculated regional morphometric similarity following a previously published protocol (Seidlitz et al., 2018), see Figure B-A and B-B. We assessed the replicability of our approach by comparing regional morphometric similarity maps to previously published maps (Morgan et al., 2019). The correlation between our regional morphometric similarity maps and the previously published maps by Morgan et al. (2019) was 0.8, thus indicating good replicability; see the Supplemental text and Supplemental Figure A for details.

### Mapping the DKT atlas regions to functional networks

We focused on seven widely recognized functional brain networks derived from an analysis of an independent resting state fMRI dataset (Thomas Yeo et al., 2011), see Supplemental Figure B. To obtain a correspondence between the regions of the DKT atlas and the seven functional networks we followed the approach by (Váša et al., 2018). Briefly, for each region of the DKT atlas we calculated the overlap (as the number of vertices) between each region of the DKT atlas and with each of the seven functional networks. Thereafter, for each region, the largest overlap determined to which functional network an region was ascribed. This resulted in a mapping of each region to one of the seven functional networks.

### Normative modeling of morphometric similarity

For normative modeling, each of the ten datasets containing only healthy controls was split into a training set (90%) and a test set (10%), see Figure A. For the clinical dataset, all individuals with schizophrenia as well as 80% of the healthy controls were included in the test set while the remaining healthy controls (20%) were included in the training set. The per-dataset splits were created using the createDataPartition function from the caret R package, preserving the distribution of age, sex, and scanner (some datasets contained multiple scanners) in both the training and the test split of each dataset. The training set included 4,310 participants and the test set 859 participants. Bayesian Linear Regression (BLR) with likelihood warping using a B-spline (spline order=3, number of knots=5) was used to predict morphometric similarity from a vector of covariates (age, sex, Euler number, and scanner) (Rutherford et al., 2023). For a complete mathematical description and explanation of this implementation, see (Fraza et al., 2021). Briefly, for each brain region of interest (r),

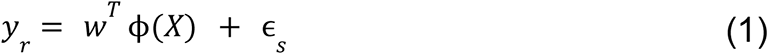

where y is the predicted distribution of morphometric similarity for region r, *w^T^* is the estimated weights vector, *X* is the set of covariates, ϕ(*X*) the B-spline basis expansion applied to them, and ɛ_*s*_ a gaussian noise distribution term for scanner *s*.

Individual deviations: infra- and supra-normal morphometric similarity

Our normative model is cross-sectional. We followed the method proposed by Barbora Rehák Bučková et al (2024) who showed that longitudinal normative modeling metrics (see Statistics) for adult individuals with schizophrenia and healthy participants can be calculated from a large cross-sectional normative model of healthy participants (Barbora Rehák Bučková et al., 2024). We created morphometric similarity normative charts for each region using the training set consisting of healthy participants. Thereafter, individuals from the test set were positioned on the morphometric similarity normative charts. For each individual deviation (z) scores, quantifying individual deviation from the normative average, were calculated for all brain regions (r) and all participants (n), as follows:

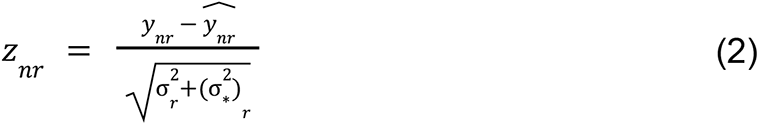

where *y*_*nr*_ means the true morphometric similarity value and 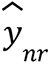 is the predicted mean morphometric similarity value. The difference in these values is normalized to account for two different sources of variation; i) 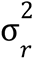, which is the aleatoric uncertainty and reflects the variation between individuals across the population, and ii) the epistemic uncertainty, 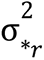, which accounts for the variance associated to modeling uncertainty introduced by the model assumptions or parameter selection. Z-scores were then categorized as either: (i) normal, i.e. within the normative range of variation for healthy individuals with the same age range, sex, Euler number, and scanner; (ii) supra-normal: significantly exceeding the normative range; or, (iii) infra-normal: significantly below the normative range. As per previous studies (Bethlehem et al., 2020; Di Biase et al., 2022; ENIGMA Clinical High Risk for Psychosis Working Group et al., 2024; Huang et al., 2024; Janssen et al., 2021; Lv et al., 2021; Rutherford et al., 2023) we considered a z-score ≧ than 1.96 as supra-normal and a z-score ≦ -1.96 as infra-normal.

### Evaluation of the normative model

The unseen data in the complete test set was used to evaluate the normative model. The evaluation included Q-Q plots, and the metrics proportion of explained variance, mean standardized log loss, standardized mean square error, root mean square error, rho, bayesian information criterion, skewness, and kurtosis (see Supplemental Figure C).

### Statistical analyses

#### The typical age-dependency of regional morphometric similarity

Normative modeling allowed us to assess the age-dependency of normative morphometric similarity across cortical regions from late childhood to old age. We tested whether the regions could be clustered into distinct groups of regions based on the shape of the age-dependence of MS, see Figure B-D. We therefore binned the training set along the age range into 27 equally-sized age bins (3.6 years/bin). Thereafter, a 62x27 matrix was constructed where each row was a region and each column contained the median normative morphometric similarity (50th percentile) of a particular age bin. Principal Component Analysis was applied to the matrix using the prcomp R package. The resulting Euclidean distance matrix was used for hierarchical clustering. The optimal number of clusters was determined using the NbClust R package, which generates 23 cluster solutions using different methods and from which the optimal number of clusters is chosen by majority voting (Charrad et al., 2014).

#### Effects of diagnosis on morphometric similarity of functional networks

Averaged z-scores of the seven functional networks were compared between healthy controls and individuals with schizophrenia at baseline using Welch t-tests. Effect sizes are given as Cohen’s d. Only p values that survived FDR correction for multiple comparisons were considered significant.

#### The percentage of individuals with infra- or supra-normal deviance per region

We calculated for each region (r) and at each timepoint (*t* = *t*1 or *t*2) the percentage of patients and healthy controls that had supra- or infra-normal z-scores:

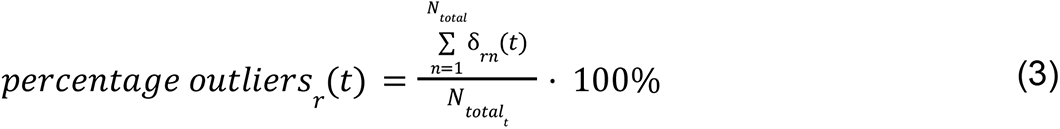

where δ_*rn*_ (*t*) equals 1 if participant *n*=1,…,*N_total_* has |Z| > 1.96 in region r at time point *t*, and equals 0 otherwise and *N_total_* is the total number of participants at time point *t*. Group differences for each region and at each time point in the proportion of individuals with supra- or infra-normal z-scores were examined using the two-proportions z-test.

Differences in percentage of outliers over time by region are calculated by subtracting the baseline percentage from the follow-up percentage for each region r:

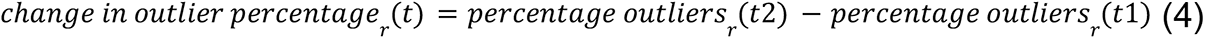

To assess the spatial distribution of infra- and supra-normal deviations we built diagnostic-wise brain maps, see Figure B-E.

#### Effects of diagnosis on the amount of outlier regions per participant

Here we determine the change over time in total number of outlier regions per participant as follows:

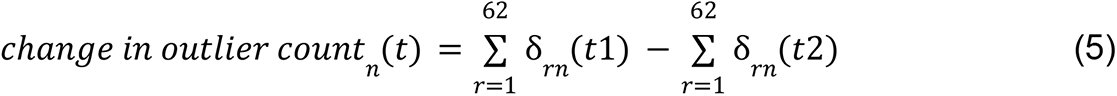

where *δ* equals 1 if participant n has |Z| > 1.96 for region r=1,…,62, and equals 0 otherwise. Diagnostic group differences in the change in total number of outlier regions by participant were examined using a Welch t-test. In the group of individuals with schizophrenia we tested for associations between change in outlier count over time and IQ and PANSS positive, negative and general scores using Pearson correlation.

### Supplemental analyses

Firstly, we repeated the analysis of clustering of regions based on normative models for males and females separately to assess whether results differed by sex. Secondly, we calculated infra- and supra-normal deviance for males and females separately to assess whether results differed by sex. Thirdly, we assessed whether using z-scores from normative modeling led to stronger diagnostic group results as compared to using ‘raw’ morphometric similarity (i.e., the traditional approach). Fourthly, we averaged morphometric similarity across all regions and hemispheres to determine whole brain morphometric similarity and compared this between cases and controls.

## Results

### Visualization and evaluation of the normative models

An exemplary normative model plot with percentile curves for the left hemispheric superior frontal gyrus can be seen in Figure B-C. Distributions of the evaluation metrics of the normative models can be found in Supplemental Figure C.

**Figure B.**
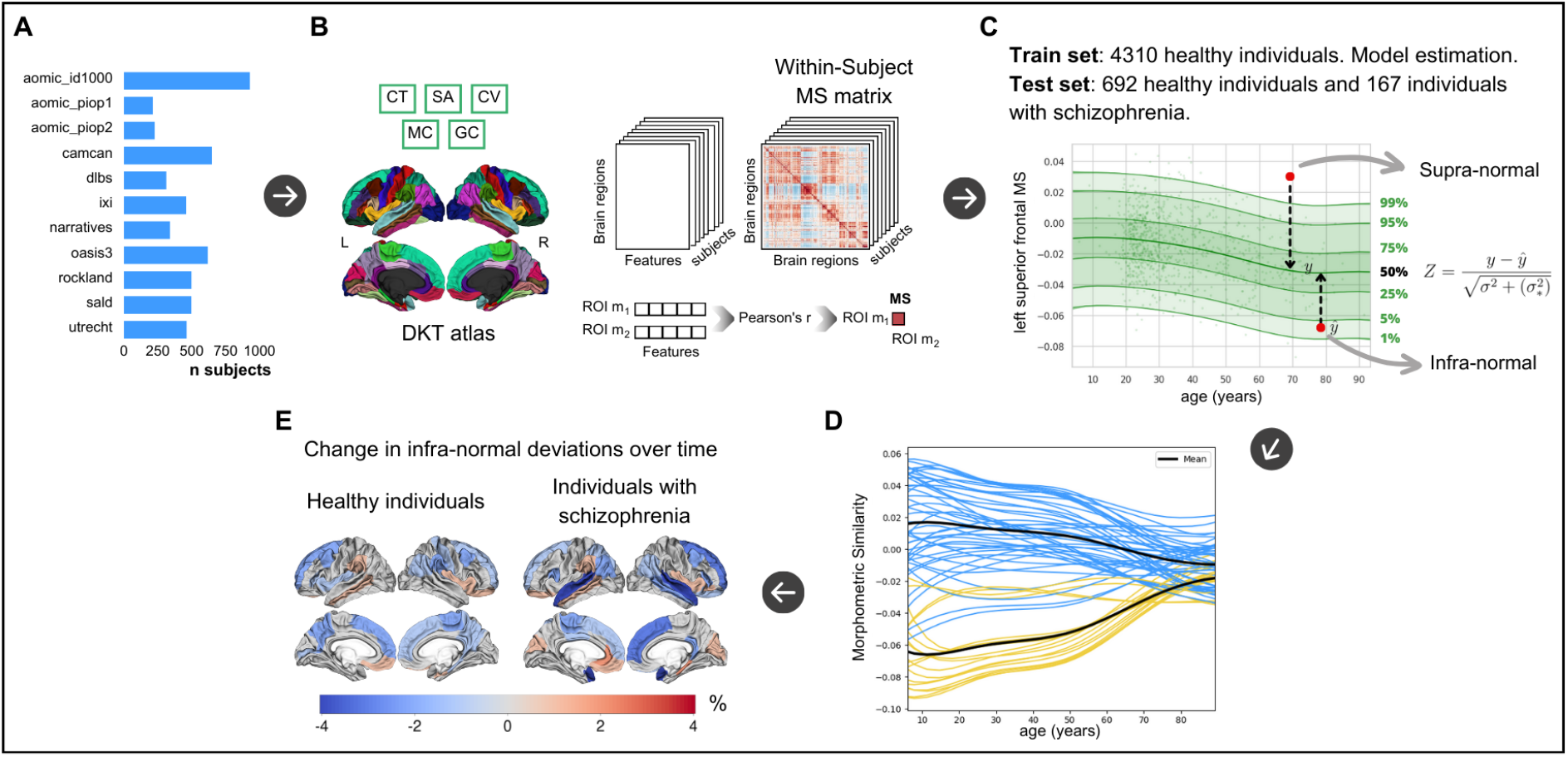
Study overview. **A**. Ten cross-sectional datasets consisting of healthy individuals only and one longitudinal dataset consisting of healthy controls and individuals with chronic schizophrenia (utrecht) were used in the study. **B**. morphometric similarity matrices were constructed following established protocols using cortical thickness, cortical volume, surface area, mean curvature and Guassian curvature extracted for each cortical region (Seidlitz et al., 2018). **C**. Example centile plot from normative modeling of regional morphometric similarity of the left hemispheric superior frontal gyrus for assessing individual deviance (Z). Normative modeling was done following established protocols (Rutherford et al., 2022) using prediction on unseen test data following training of the model. **D**. Normative models of each cortical region were clustered based on their typical age-dependency. **E**. Cortical maps depicting percentage longitudinal change of extreme deviance below the norm, i.e. infra-normal deviance, in individuals with schizophrenia and healthy individuals. n, number of subjects, CT, cortical thickness; SA, surface area; CV, cortical volume; MC, mean curvature; GC, Gaussian curvature. MS, morphometric similarity.

### The typical age-dependency of regional morphometric similarity

Principal component analysis using the age-dependency of regional morphometric similarity revealed that the first two principal components explained 98.63% of the variance (first component: R^2^=77.14%, second component: R^2^=21.49%) of the median normative morphometric similarity across age groups and were therefore selected for constructing the distance matrix, see Figure C. Clustering gave two clusters as the optimal solution. As can be seen in Figure C, clustering was extremely similar between left and right hemispheres. The first cluster (number of regions = 44, 71% of all regions) includes mostly frontal and temporal regions with positive morphometric similarity tending towards zero during aging. The second cluster (18 regions, 29%) consists of inferior prefrontal, postcentral gyrus and occipital regions with negative morphometric similarity tending towards zero during aging.

39% of all regions from Cluster 1 were part of the default mode network which is significantly more than expected (P<0.001, one sample z-test for proportions, see Table B). For Cluster 2 this was 11% (P=0.48).

**Figure C.**
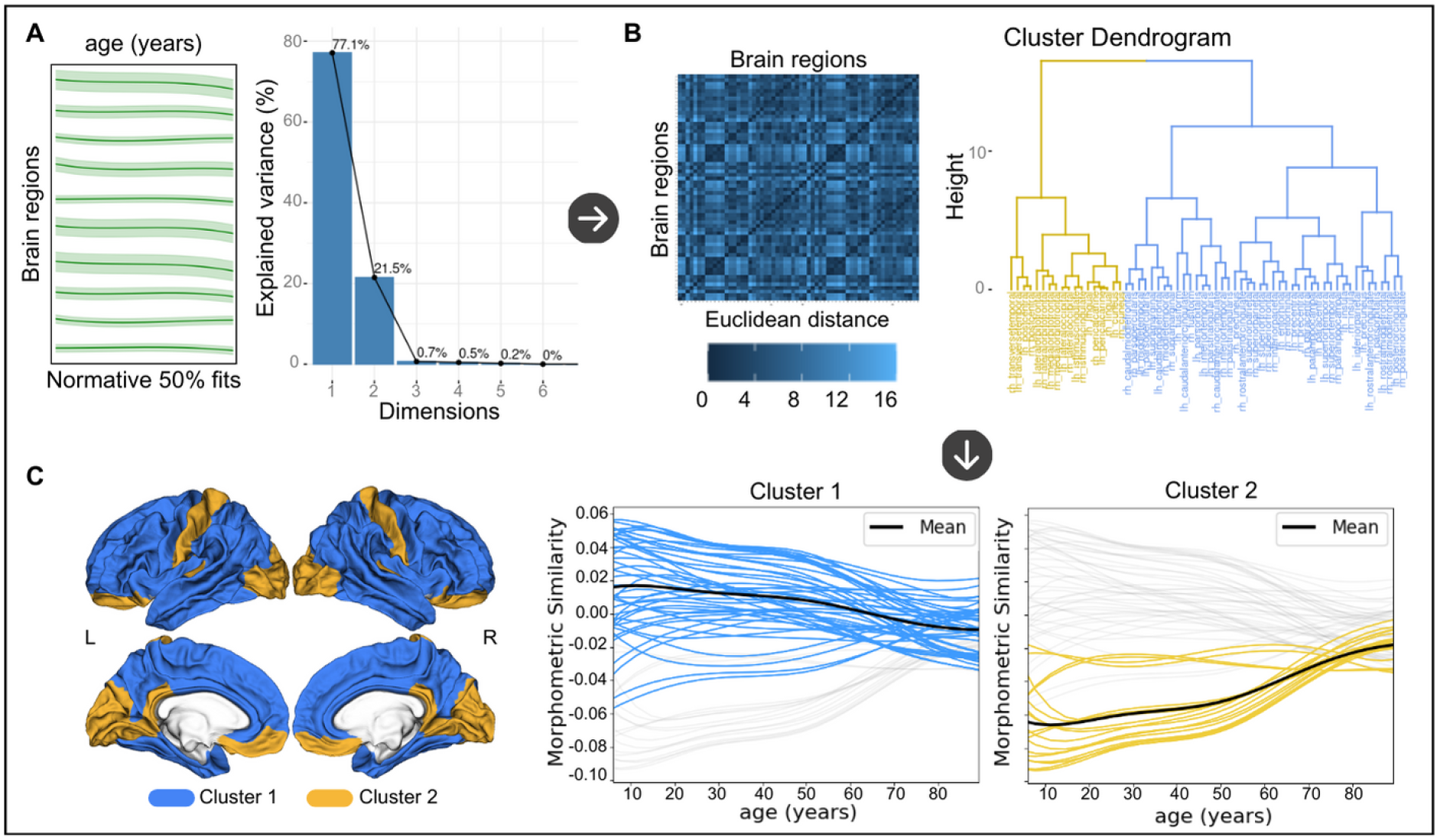
Clustering of brain regions based on the typical age-dependency of regional morphometric similarity. **A**. The median normative morphometric similarity of the normative model of each cortical region was entered into a Principal Component Analysis (PCA). The first two components explaining 98.6% of the variance were selected. **B**. The Euclidean distance matrix was extracted from the PCA output and used for hierarchical clustering. A two cluster solution was determined optimal. **C**. Each plot shows the median normative fits for a cluster with the remaining normative fits in gray. The average fit across all fits belonging to a cluster is added in black. The cluster solution was also visualized on the cortex displaying the distribution of the two clusters across the cortex. L, left hemisphere, R, right hemisphere. The upper row are lateral views, the bottom row are medial views.

**Table B.** Number of regions from each of the two clusters belonging to each of the seven functional networks. MS, morphometric similarity.

| Network | MS<br>Cluster 1 | MS<br>Cluster 2 |
| --- | --- | --- |
| Visual | 2 | 6 |
| Somato Motor | 2 | 2 |
| Dorsal Attention | 4 | 4 |
| Ventral Attention | 8 | 0 |
| Limbic | 9 | 0 |
| Fronto Parietal | 2 | 4 |
| Default Mode | 17 | 2 |

### Effects of diagnosis on morphometric similarity of functional networks

At baseline, the group of individuals with schizophrenia had a positive average deviance of morphometric similarity for the somatosensory network which differed significantly from the negative average deviance of morphometric similarity in healthy controls (d=0.30; p<0.01). Individuals with schizophrenia had a negative average deviance of morphometric similarity for the default mode network which differed significantly from the positive average deviance of morphometric similarity in healthy controls (d=-0.36; p<0.001), see Table C and Figure D. At follow-up, the group of individuals with schizophrenia maintained the negative morphometric similarity for the default mode network compared to healthy controls but this difference did not withstand correction for multiple comparisons, see Figure D.

**Table C.** Results of statistical tests for case-control differences in z-scores averaged across regions belonging to the seven functional brain networks (Thomas Yeo et al., 2011). A positive t-statistic indicates that the group of individuals with schizophrenia has a higher mean than the group of healthy controls, while a negative t-statistic indicates that the group of individuals with schizophrenia has a lower mean MS than the group of healthy controls. *P_FDR_ < 0.05.

| <u>Timepoint</u> | <u>Network</u> | <u>z-scores</u> |  |
| --- | --- | --- | --- |
|  |  | *t-*<br>*statistic* | *P value*<sup>A</sup> |
| Baseline | Visual | 1.451 | 0.148 |
|  | Somato<br>Motor | 2.940 | 0.003* |
|  | Dorsal<br>Attention | -0.844 | 0.399 |
|  | Ventral<br>Attention | -0.075 | 0.940 |
|  | Limbic | -1.902 | 0.058 |
|  | Fronto<br>Parietal | -0.805 | 0.421 |
|  | Default<br>Mode | -3.665 | 0.0002<br>* |
| Follow-up | Visual | 1.519 | 0.130 |
|  | Somato<br>Motor | 1.675 | 0.095 |
|  | Dorsal<br>Attention | -0.867 | 0.387 |
|  | Ventral<br>Attention | -0.819 | 0.413 |
|  | Limbic | -0.617 | 0.537 |
|  | Fronto<br>Parietal | -0.361 | 0.718 |
|  | Default<br>Mode | -2.046 | 0.041 |

**Figure D.**
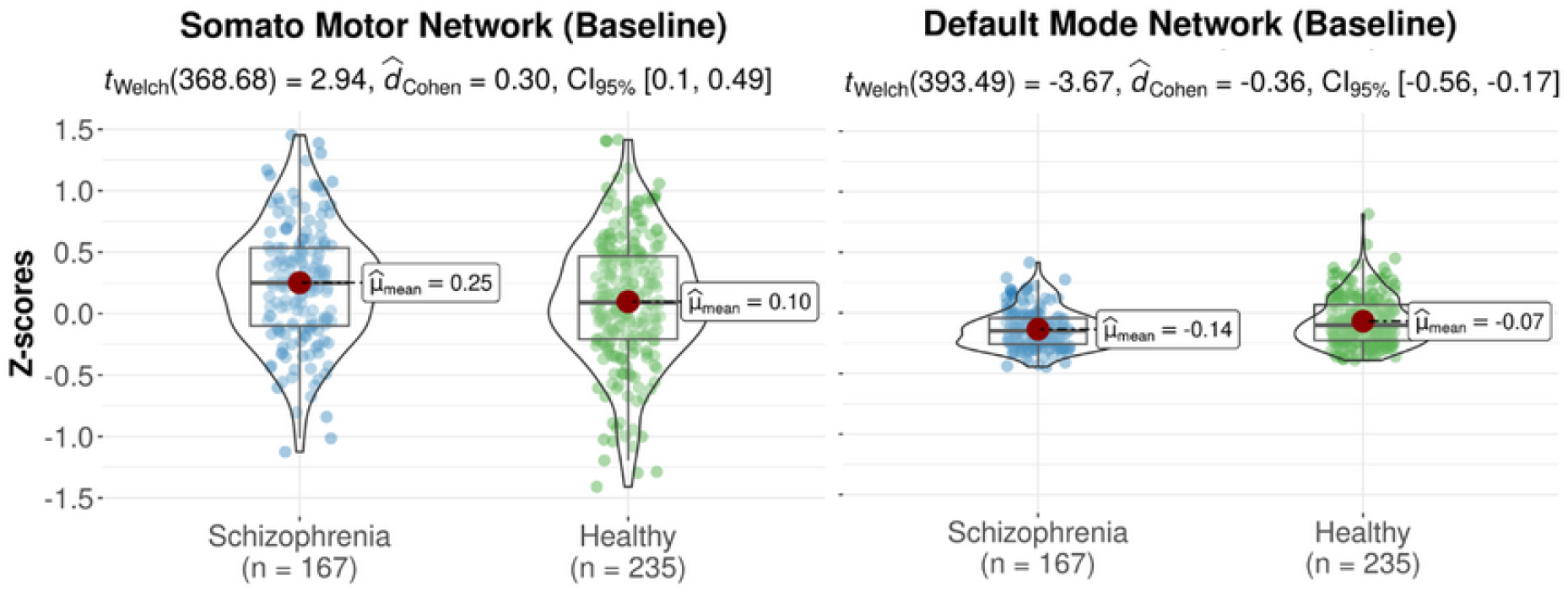
Violin plots of significant (FDR-corrected) differences in z-scores of morphometric similarity of two functional networks that differed significantly between the group of healthy controls and the group of individuals with schizophrenia at baseline.

### The percentage of individuals with infra- or supra-normal deviance per region

The percentage of individuals from the test set with either infra- or supra-normal deviance was below 6% for both the healthy individual and the schizophrenia samples at the baseline and follow-up visits (see Figure E-A and E-B). At both timepoints, the percentage of infra-normal regional morphometric similarity z-scores ranged between 0 and 6.0% for individuals with schizophrenia and between 0 and 4.3% for healthy individuals; for supra-normal morphometric similarity z-scores percentage ranges were 0–5.4% and 0–6.0%, respectively. There were no significant differences between patients and controls in the percentage of participants with infra- or supra-normal regional values at either baseline or follow-up (P>0.05). At baseline and follow-up, the percentages of individuals with schizophrenia with infra-normal deviance were highest in the superior temporal and superior frontal regions. For both diagnostic groups, the percentages of participants with supra-normal deviance were highest in the occipital lobe and postcentral gyrus (see Figure E-A and E-B).

We then assessed differences in longitudinal change in the percentage of outlier individuals for each region between cases and controls, separately for infra- and supra-normal deviance. For a majority of regions, and both for cases and controls, the change in percentage of outliers with infra-normal deviance was negative while this was not the case for the change in percentage of supra-normal deviance (see Figure E-C). This indicated that for the majority of regions, ‘normalizing’ of outlier participants over time occurred more frequently in regions where those outliers initially had infra-normal deviance compared to regions where those outliers initially had supra-normal deviance. For infra-normal deviance the regions with highest percentages of change were the bilateral superior temporal and left superior and medial frontal regions with a decrease over time of 4.2%, 0.6% and 1.8%, respectively. For supra-normal deviance the region with the highest percentage of change was the left postcentral gyrus with an increase of 1.2% (see Figure E-C).

**Figure E.**
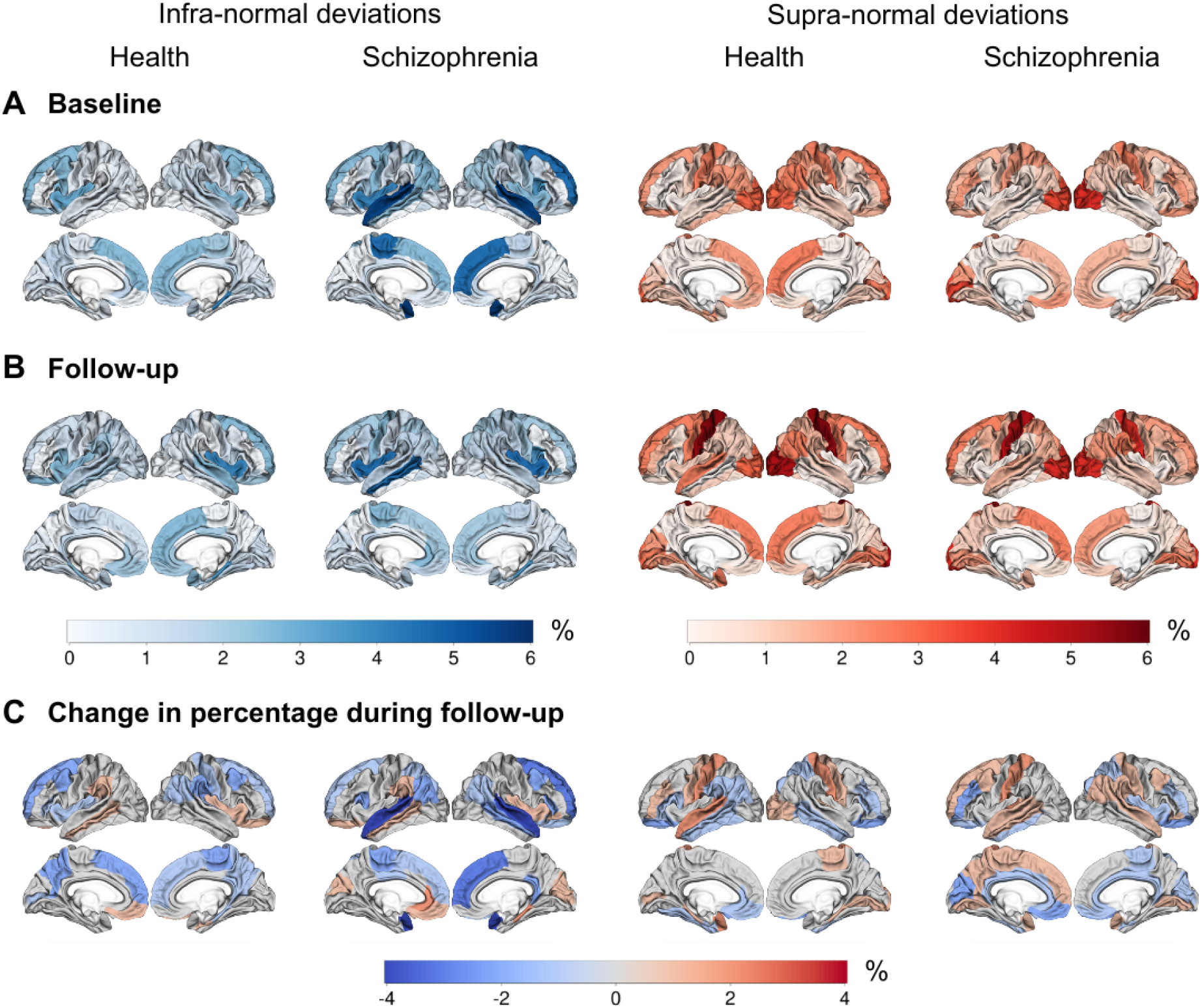
Cortical maps showing the percentage of individuals with infra- and supra-deviance morphometric similarity per region, **A**) at baseline, **B**) at follow-up, and **C**) the change in percentage during follow-up (i.e., percentage at follow-up - percentage at baseline). A negative percentage means that the regional percentage of individuals with infra- or supra-normal deviance decreased during follow-up.

### Effects of diagnosis on the amount of outlier regions per participant

No significant group differences for the average cross-sectional total outlier count and change in total outlier region count over time were found between healthy controls and individuals with schizophrenia (see Supplemental Figure D).

In the group of individuals with schizophrenia no significant correlations were found between cross-sectional outlier region count or change in outlier region count over time and IQ and PANSS scores (P>0.05).

### Supplemental analyses

Clustering of regions based on normative models showed three clusters in males and two clusters in females as the optimal solution (see Supplemental Figure E).

Across both time points and regions, the percentage of female participants with either infra- or supra-normal deviance at baseline and follow-up was higher compared to males. In both sexes there were no significant differences in the percentage of individuals with infra- or supra-normal regional values at baseline and follow-up (see Supplemental Text and Supplemental Figures F and G).

We replicated the finding of lower average deviance morphometric similarity for the default mode network for the group of individuals with schizophrenia compared to healthy controls in the analyses using raw values at baseline, albeit with a smaller effect size (d=-0.27, p<0.01) compared to the normative-modeling based z-scores (see Supplemental Table A, Supplemental Text and Supplemental Figure H).

There was no difference of whole brain morphometric similarity between individuals with schizophrenia and healthy individuals (see Supplemental Figure I).

## Discussion

To the best of our knowledge this study is the first longitudinal study of morphometric similarity, as well as the first to assess morphometric similarity in a normative modeling framework. We established regional age-dependent normative trajectories for morphometric similarity across the adult age range. We found that, overall,mean regional morphometric similarity converged towards zero with increasing age, which is coherent with a prior study (Zhukovsky et al., 2022). Here, we extended prior findings by showing that age-dependent trajectories can be clustered into two groups. One cluster contains trajectories that start with positive morphometric similarity values at younger ages which decrease to zero with age. The other cluster starts with negative morphometric similarity values which increase to zero with age. We also showed that the default mode network contained disproportionally more regions of the former compared to the latter cluster. Studies have consistently reported age-related reductions in resting-state functional connectivity of the default mode network as well as structural network connectivity (Cox et al., 2016; Geerligs et al., 2015). As such, these findings are in line with the observed decline of morphometric similarity across a large age range. In addition, this finding provides support for the notion that brain structure, at least partially, underlies function (Pang et al., 2023; Zhukovsky et al., 2022).

Our investigation aimed to compare morphometric similarity across functional networks and regions between individuals with schizophrenia and healthy subjects over time.

Our cross-sectional findings revealed that, at baseline, individuals diagnosed with schizophrenia exhibited increased morphometric similarity across regions pertaining to the somato-motor network and decreased morphometric similarity across regions pertaining to the default mode network when compared to the healthy control subjects. The reduced morphometric similarity in the group of individuals with chronic schizophrenia aligns with previous research morphometric similarity findings as well as general default mode dysfunction in high-risk, early-onset and chronic schizophrenia (Morgan et al., 2019; Whitfield-Gabrieli et al., 2009; Yao et al., 2023). A recent study demonstrated that morphometric similarity was positively correlated with the likelihood of cellular network structures forming axonal connections (Lin et al., 2024). In schizophrenia, axonal dysconnectivity between networks might lead to the micro-structural and functional connectivity deficits observed across various age groups affected by the disease, including children, adolescents, and adults (Barth et al., 2023; Heuvel and Sporns, 2019; Kelly et al., 2018).

The increased morphometric similarity observed in the somatomotor network poses a complex challenge for explanation, considering the general neglect of the somatomotor network within psychiatric models of schizophrenia. Speculatively, it may be that increased morphometric similarity for the somatomotor network may be compensatory as individuals with schizophrenia may be less able to draw on other networks. Nonetheless, a recent resting-state fMRI study shed light on altered somatomotor network connectivity, suggesting its potential relevance across various psychiatric disorders, associating it with psychopathology, cognitive impairment, and impulsivity (Kebets et al., 2019).

In a minority of individuals with schizophrenia, infra-normal and supra-normal deviations of morphometric similarity were present (<6%), mirroring observations akin to those found in healthy individuals. Notably, outlier percentages did not differ significantly between baseline and follow-up assessments. These results indicate that variations in morphometric similarity among individuals with schizophrenia predominantly existed within the spectrum of variations observed in the healthy reference sample (Bedford et al., 2023; ENIGMA Clinical High Risk for Psychosis Working Group et al., 2024; Segal et al., 2023; Winter et al., 2022; Wolfers et al., 2018). Previous normative modeling studies conducted on individuals with schizophrenia or those at high risk for the condition have reported similar percentages for cortical volume and cortical thickness (ENIGMA Clinical High Risk for Psychosis Working Group et al., 2024; Segal et al., 2023; Wolfers et al., 2018). We demonstrated significant heterogeneity in morphometric similarity among individuals with schizophrenia, suggesting that structural deficits in schizophrenia are nuanced and intricate, which contributes to a substantial overlap between individuals with chronic schizophrenia and healthy controls.

The change in total outlier count over time did not differ between cases and controls, meaning that individuals with schizophrenia did not have an accelerated accumulation of outlier regions as compared to the healthy individuals during follow-up. In individuals with Alzheimer’s disease, the total outlier count for cortical thickness was increased at baseline and also progressed over time when compared to healthy individuals implying that the outlier count can be used to track neurodegeneration in Alzheimer’s disease (Verdi et al., 2023). While morphometric similarity is being increasingly used to detect macroscopic brain abnormalities in individuals with psychiatric disorders, with both regional increases and decreases reported, there is no prior information available about changes of morphometric similarity over time (Li et al., 2021; Morgan et al., 2019). In addition to the change in outlier count we showed that the increase and decrease of the regional outlier percentage during follow-up was relatively small and similar in both cases and controls (<|4|%). This indicates that of those with baseline regional morphometric similarity values outside the normative range few fluctuated considerably over time. For schizophrenia it may mean that change in regional cortical thickness over time, as assessed by normative modeling, is more sensitive to disease effects compared to regional morphometric similarity (Barbora Rehák Bučková et al., 2024). Taken together, our results could imply that morphometric similarity may have restricted utility in elucidating the pathophysiology of schizophrenia (Winter et al., 2022).

Normative models may be better able to detect diagnosis-related effects compared to raw, i.e. traditional, data models because normative modeling allows for the consideration of numerous sources of variance. Some of these sources may not carry clinical significance (e.g., scanner variability, Euler number), while others simultaneously encapsulate clinically relevant information within the framework of a reference cohort. Our supplemental results are in line with this, i.e. our study demonstrated increased statistical significance using normative models when compared to raw data models. Thus, normative modeling may possess the capacity to capture overarching population patterns, discern clinical disparities between groups, and retain the ability to investigate individual differences (Rutherford et al., 2023).

This study has limitations. For clinical translation, progressions in normative modeling stand as a crucial prerequisite, underscoring the necessity for a broadened diversification of datasets. While the present study benefited from a substantial training set, larger and more heterogeneous datasets, incorporating not solely European ancestry data, are imperative for heightened performance and validity (Bethlehem et al., 2022). The normative modeling framework employed herein treats each region as independent, notwithstanding potential correlations among z-scores from neighboring regions. To mitigate this challenge, one plausible strategy involves implementing data reduction techniques, such as principal component analysis applied to the z-scores (Rutherford et al., 2022). While our sample size of individuals with schizophrenia was comparable to previous normative modeling studies, larger (multicenter) clinical samples may enable the identification of distinct clusters of patients exhibiting significant deviance (Rutherford et al., 2023; Wolfers et al., 2018). In conclusion, the current study used normative modeling to show decreased morphometric similarity of the default mode network in a group of individuals with chronic schizophrenia, replicating previous findings in groups of individuals with first-episode psychosis. Using a longitudinal design we showed that the change in total number of outlier regions over time was not different between cases and controls. Change over time of the regional percentage of outliers was low indicating only few participants showed large changes over time. Normative modeling demonstrated that significant cross-sectional reductions and longitudinal changes of morphometric similarity are only present in a minority of individuals with schizophrenia. Our study provides a lay out for future studies using normative modeling including cross-sectional and longitudinal neuroimaging phenotypes.

## Data availability

The study IDs of the included participants from the ten publicly available datasets, the five regional metrics and regional morphometric similarity from the ten publicly available datasets, the overlap between the DKT atlas and the 7 functional networks, the normative model plots with percentile curves, the Q-Q plots, the information and usage instructions about the docker we created for normative modeling are all available at https://github.com/iamjoostjanssen/NormModel_MorphoSim_SZ.

Aomic (id1000, piop1 and piop2) is available at https://openneuro.org/datasets/ds003097, https://openneuro.org/datasets/ds002785 and https://openneuro.org/datasets/ds002790, camcan is available at https://camcan-archive.mrc-cbu.cam.ac.uk, dlbs is available at https://fcon_1000.projects.nitrc.org/indi/retro/dlbs, ixi is available at http://brain-development.org/ixidataset, narratives is available at https://openneuro.org/datasets/ds002345, oasis3 is available at www.oasis-brains.org, rockland is available at http://fcon_1000.projects.nitrc.org/indi/enhanced and sald is available at http://fcon_1000.projects.nitrc.org/indi/retro/sald.

## Acknowledgements

Supported by the Spanish Ministry of Science and Innovation, Instituto de Salud Carlos III (ISCIII), CIBER-Consorcio Centro de Investigación Biomédica en Red- (CB/07/09/0023), co-financed by the European Union, ERDF Funds from the European Commission, “A way of making Europe”, (PI16/02012, PI17/01249, PI17/00997, PI19/01024, PI20/00721, PI22/01824, PI22/01621, PI23/00625), financed by the European Union - NextGenerationEU (PMP21/00051), Madrid Regional Government (S2022/BMD-7216 AGES 3-CM), European Union Seventh Framework Program, European Union H2020 Program under the Innovative Medicines Initiative 2 Joint Undertaking: Project PRISM-2 (Grant agreement No.101034377), Project AIMS-2-TRIALS (Grant agreement No 777394), Project COllaborative Network for European Clinical Trials For Children “c4c” (Grant agreement No 777389) Horizon Europe, the National Institute of Mental Health of the National Institutes of Health under Award Number 1U01MH124639-01 (Project ProNET), Award Number 5P50MH115846-03 (Project FEP-CAUSAL) and Award Number 1R01MH128971-01A1 (Project SZ-aging), Fundación Familia Alonso, and Fundación Alicia Koplowitz. The authors thank Yasser Alemán-Goméz, Alberto Fernández Pena, Zimbo Boudewijns, and Joyce van Baaren for code and technical assistance.

