## Supplement for "Heterogeneity of morphometric similarity networks in health and schizophrenia"

**Content**

**Supplemental Table**

Supplemental Table A: ‘Raw’ morphometric similarity 2

**Supplemental Text and Figures**

Supplemental Text and Figure A: Replicability of morphometric similarity 4

Supplemental Figure B: Seven functional networks 6

Supplemental Figure C: Performance metrics of normative models 7

Supplemental Figure D: Effects of diagnosis on the amount of outlier regions per participant 8

Supplemental Figure E: Clustering of brain regions based on normative models of morphometric similarity per sex 9

Supplemental Figure F: Infra- and supra-normal deviances in males 10

Supplemental Figure G: Infra- and supra-normal deviances in females 12

Supplemental Figure H: ‘Raw’ morphometric similarity 14

Supplemental Figure I: Global morphometric similarity 16

**References** 17

| Timepoint | Network | z-scores | | Mean MS | | | |
|---|---|---|---|---|---|---|---|
| | | $t$-statistic | $P\ value^A$ | $mean\ (SD)_H$ | $mean\ (SD)_{SZ}$ | $t$-statistic | $P\ value^A$ |
| Baseline | Visual | 1.451 | 0.148 | -0.021 (0.007) | -0.02 (0.007) | 1.764 | 0.022 |
| | Somato Motor | 2.940 | 0.003* | -0.015 (0.018) | -0.012 (0.017) | 1.011 | 0.313 |
| | Dorsal Attention | -0.844 | 0.399 | -0.018 (0.009) | -0.018 (0.009) | -0.199 | 0.842 |
| | Ventral Attention | -0.075 | 0.940 | -0.017 (0.007) | -0.017 (0.008) | 1.162 | 0.246 |
| | Limbic | -1.902 | 0.058 | -0.013 (0.01) | -0.014 (0.011) | -0.778 | 0.437 |
| | Fronto Parietal | -0.805 | 0.421 | -0.005 (0.013) | -0.006 (0.013) | -1.461 | 0.145 |
| | Default Mode | -3.665 | 0.0002 * | -0.008 (0.007) | -0.01 (0.006) | -2.753 | 0.006* |
| Follow-up | Visual | 1.519 | 0.130 | -0.022 (0.007) | -0.02 (0.007) | 1.721 | 0.216 |
| | Somato Motor | 1.675 | 0.095 | -0.018 (0.018) | -0.017 (0.018) | 0.461 | 0.645 |
| | Dorsal Attention | -0.867 | 0.387 | -0.017 (0.009) | -0.017 (0.009) | -0.322 | 0.748 |
| | Ventral Attention | -0.819 | 0.413 | -0.016 (0.008) | -0.017 (0.007) | -1.080 | 0.281 |
| | Limbic | -0.617 | 0.537 | -0.012 (0.011) | -0.012 (0.011) | 0.256 | 0.798 |
| | Fronto Parietal | -0.361 | 0.718 | -0.006 (0.015) | -0.007 (0.013) | -0.823 | 0.411 |
| | Default Mode | -2.046 | 0.041 | -0.008 (0.007) | -0.009 (0.006) | -1.495 | 0.136 |

Supplemental Table A. Results of statistical tests for case-control differences in z-scores and ‘raw’ (i.e. traditional values rather than normative modeling-based z-scores) morphometric similarity averaged across regions belonging to the seven functional brain networks as identified by Yeo et al. (2011) (Thomas Yeo et al., 2011). A positive t-statistic indicates that the group of individuals with schizophrenia has a higher mean than the group of healthy controls, while a negative t-statistic indicates that the group of individuals with schizophrenia has a lower mean than the group of healthy controls. *$P_{FDR} < 0.05$

For all healthy controls of the clinical sample at baseline we calculated the average morphometric similarity for all regions of the atlas that was used in Morgan et al. (2019). We then used a spatial permutation test, the spin test, to correlate our 5-features morphometric similarity map with the publicly available 7-features map from Morgan et al. (2019) (Morgan et al., 2019; Váša et al., 2018). The spin-test provides a P-value derived from a spherical permutation ($P_{perm}$), obtained by comparing the empirical Spearman's $\rho$ to a null distribution of 10,000 Spearman correlations, between our 5-features map and the randomly rotated projections of the map by Morgan et al. (2019) (Váša et al., 2018). The empirical Spearman's $\rho$ for the correlation between our 5-features map and the 7-features morphometric similarity map by Morgan et al. (2019) for healthy controls was 0.8 ($P_{perm}$<0.001) indicating good correspondence between our 5-features map and the 7-features map from (Morgan et al., 2019) (see the 5- and 7-features maps at Supplemental Figure B-A and B-B). To address the effect of scanner harmonization on replicability we repeated the replicability-analysis using COMBAT-corrected input (see Supplemental Figure B-C) (Fortin et al., 2018).

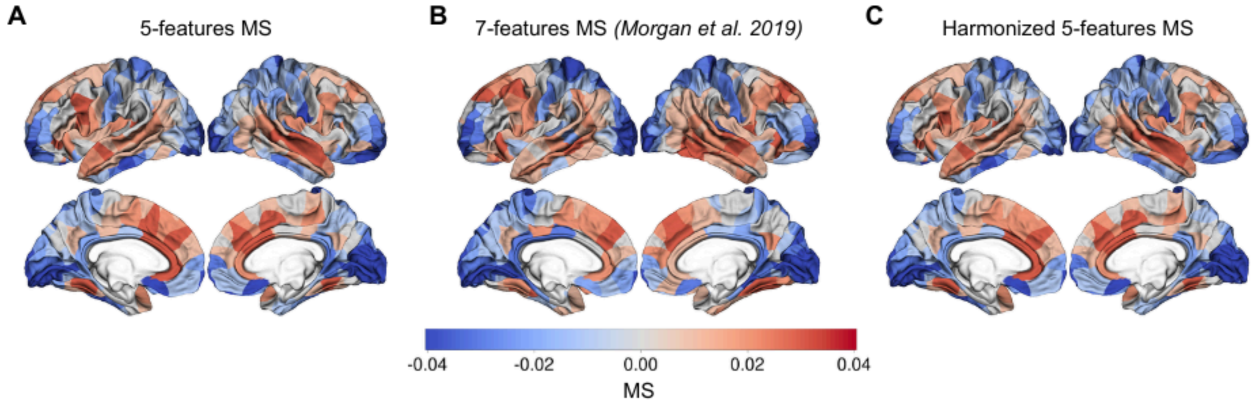

Supplemental Figure A. Replicability of MS. **A**) Reduced 5 feature approach to construct MS. **B**) morphometric similarity constructed with 7 features derived from T1-weighted and DTI (Morgan et al., 2019). **C**) Reduced 5 feature approach to construct MS, input is harmonized for use of multiple scanners before calculating MS. Scanner harmonization is done using the COMBAT method (Fortin et al., 2018). MS, morphometric similarity. As can be seen in Supplemental Figure B, the harmonized 5-features map was highly similar to the 5-features map used in the current study and the 7-features map from (Morgan et al., 2019).

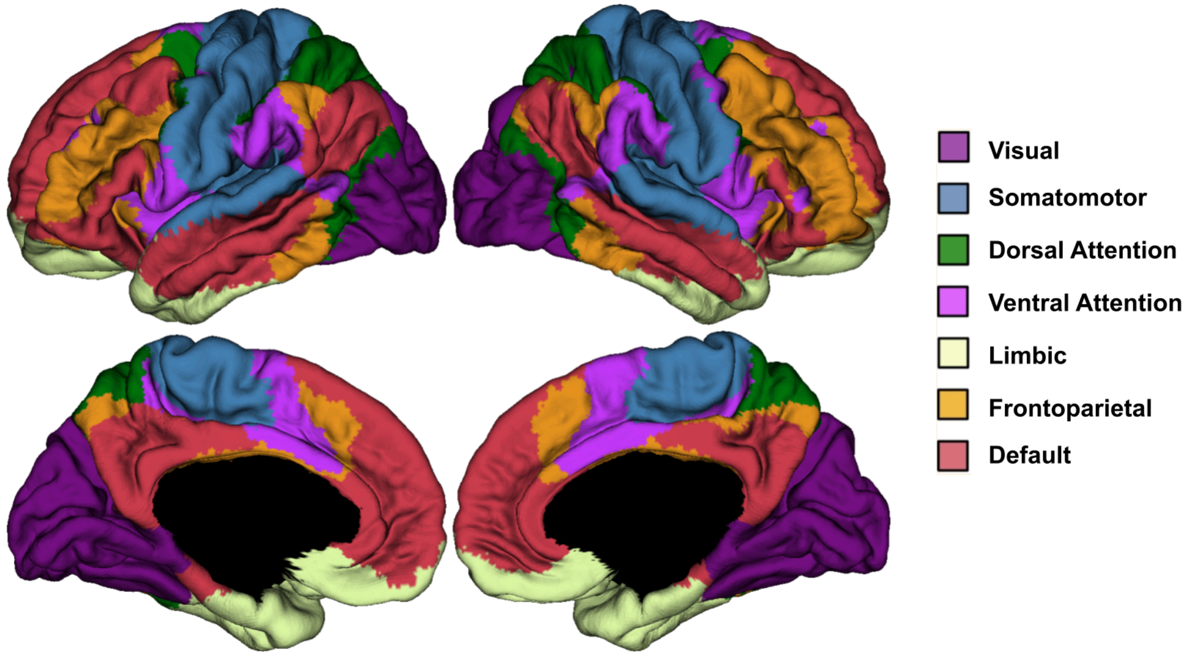


Supplementary Figure B displays the seven functional networks as published by Yeo et al. (2011).

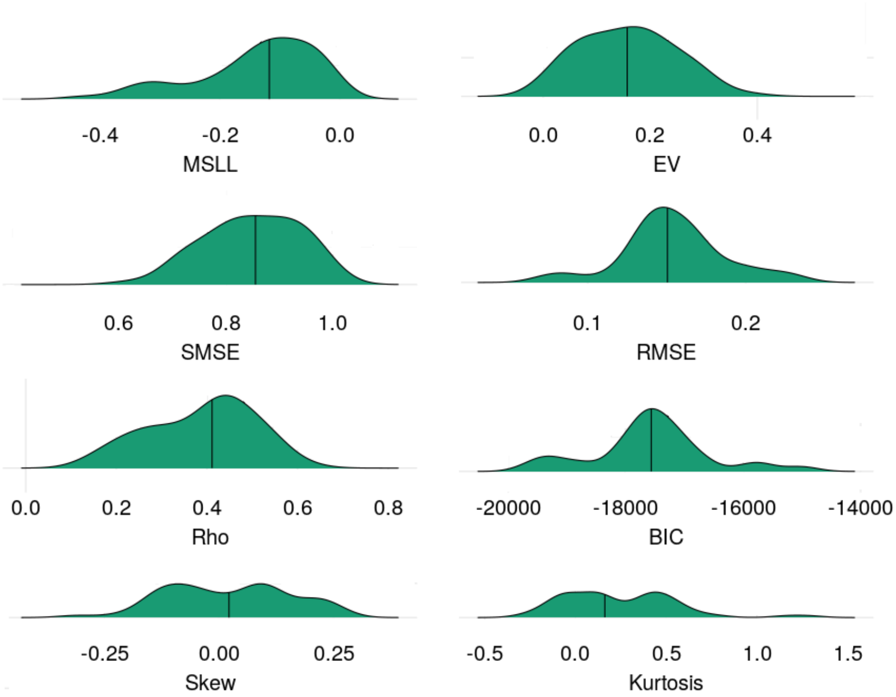


Supplementary Figure C displays the distribution and median for the performance metrics for the normative models, assessing their accuracy in estimating the relationship between morphometric similarity and age, sex, Euler number and scanner. These metrics encompass i) explained variance (EV, higher values indicating better performance), ii) Mean standardized log-loss (MSLL, lower values indicating better performance), iii) Standardized mean squared error (SMSE, lower values indicating better performance), iii) Root mean squared error (RMSE, lower values indicating better performance), Bayesian information criterion (BIC, lower values indicating better performance), iiii) Skewness (Skew), and iv) Kurtosis. Q-Q plots for all normative models are available from https://github.com/iamjoostjanssen/NormModel_MorphoSim_SZ.

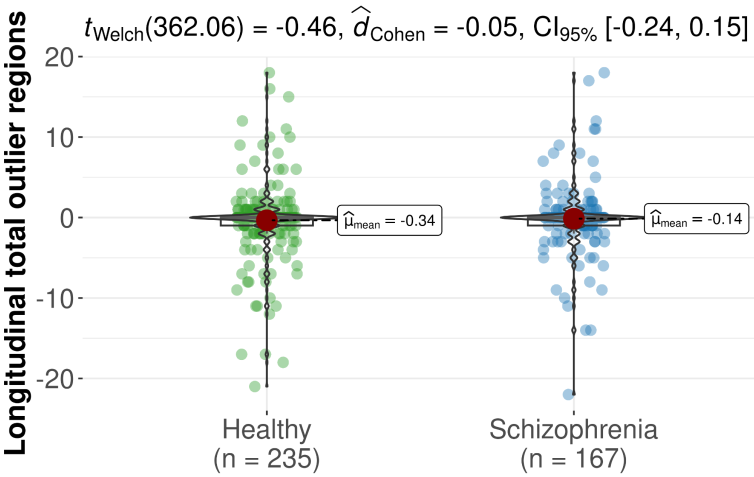


Supplemental Figure D. Longitudinal change in total outlier regions per subject (baseline - follow-up). Violin plots showing differences in longitudinal total outlier regions between the group of healthy participants and a group of individuals with schizophrenia. Effect size is given as Cohen's d. CI, 95% confidence interval. No significant diagnostic differences were found for longitudinal change in total outlier regions per subject (P=0.64). No significant correlations were found between longitudinal total outlier count and IQ and PANSS scores (P>0.05).

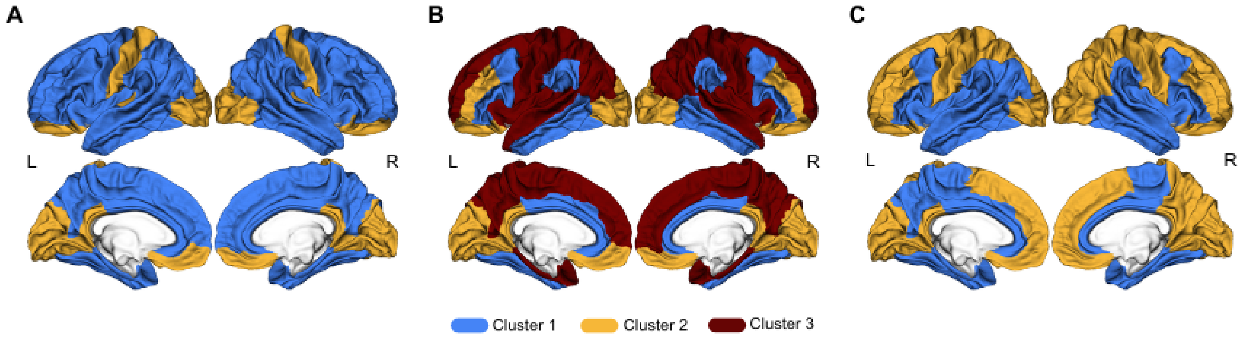


Supplemental Figure E. Cluster solution mapped to the cortex displaying the distribution of clusters across the cortex. L, left hemisphere, R, right hemisphere. The upper row are lateral views, the bottom row are medial views. Clustering for **A**) both sexes, **B**) males, **C**) females. Three clusters were found as the optimal solution for males, and two clusters for females.

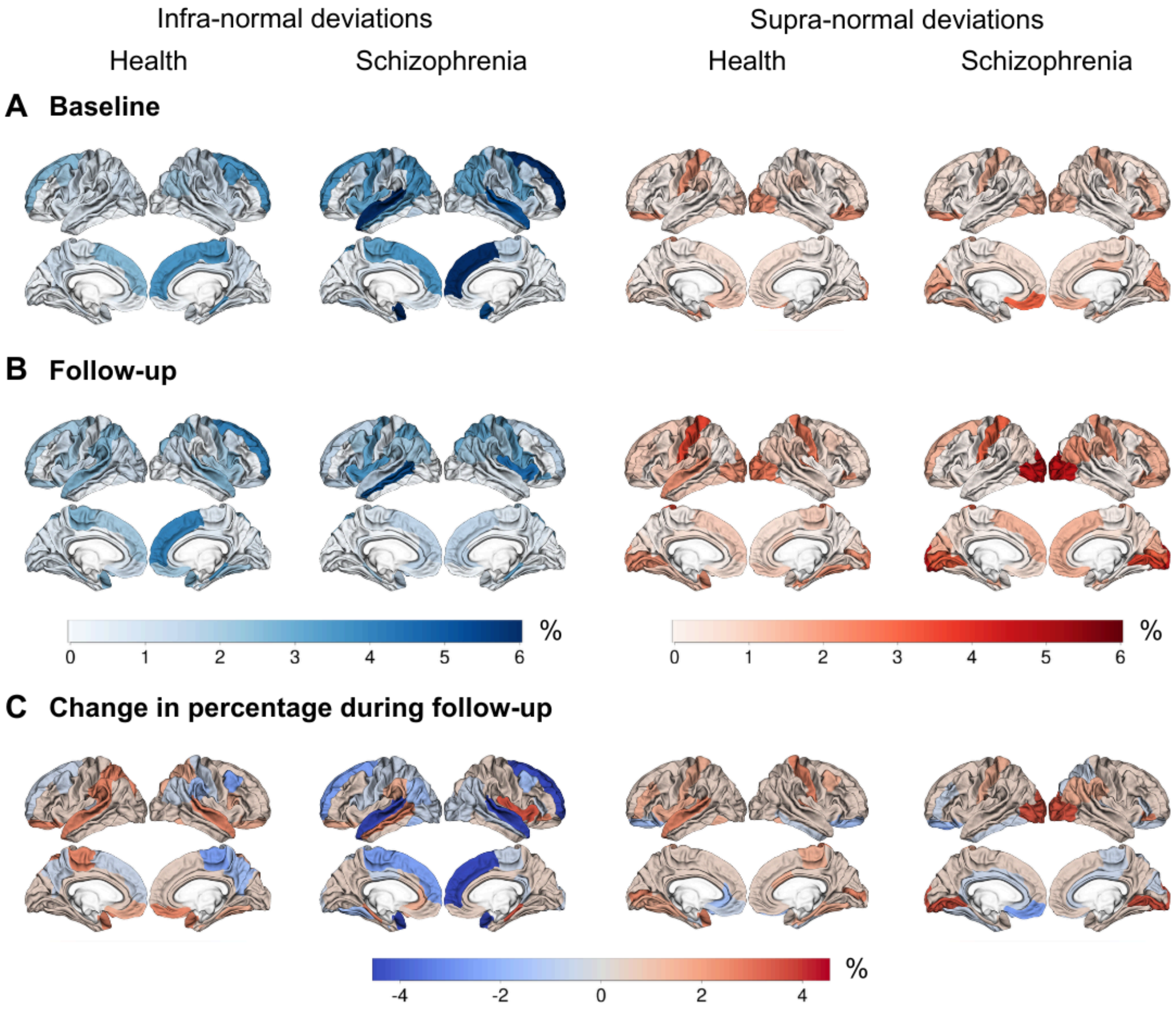


Supplemental Figure F. Cortical maps showing the percentage of *male* individuals with infra- and supra-normal deviance at **A**) baseline and **B**) follow-up and **C**) change over time (follow-up - baseline), meaning that blue indicates a decrease in the percentage of infra- and supra-normals. Across both timepoints and regions, for male individuals with schizophrenia the percentage infra-normal regional Z-scores ranged between 0 and 6.2% and for male healthy controls it ranged between 0 and 4.4%; the corresponding range for the percentages of supra-normal Z-scores were 0–5.4% and 0–4.4%, respectively, see Supplementary Figure G-A and G-B. There were no significant diagnostic differences in the percentage of male participants with infra- or supra-normal regional values at baseline and follow-up ($P>0.05$). For infra-normal deviance the regions with highest percentages of change in individuals with schizophrenia were the bilateral superior temporal and and right superior frontal regions with a decrease over time of 4.7%. For supra-normal deviance the regions with the highest percentage of change in individuals with schizophrenia were the bilateral occipital regions with an increase over time of 3.1% (see Figure G-C).

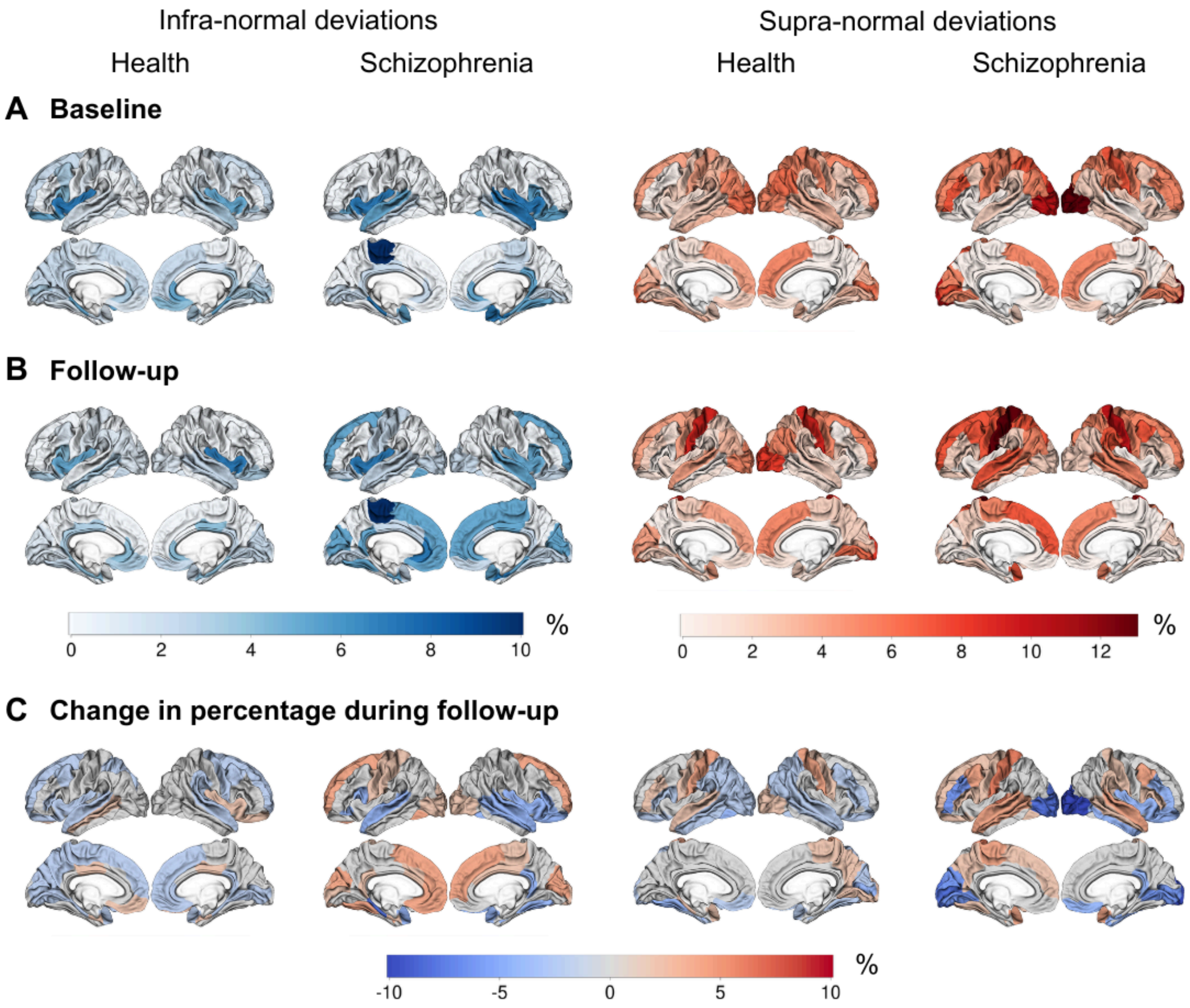


Supplemental Figure G. Cortical maps showing the percentage of fe*male* individuals with infra- and supra-normal deviance at **A**) baseline and **B**) follow-up and **C**) change over time (follow-up - baseline), meaning that blue indicates a decrease in the percentage of female infra- and supra-normals. Across both timepoints and regions, the percentage of female participants with either infra- or supra-normal deviance at baseline and follow-up was higher compared to males (see Supplementary Figure G-A and G-B). Across both timepoints and regions, for female individuals with schizophrenia the percentage infra-normal regional Z-scores ranged between 0 and 10.5% and for female healthy controls it ranged between 0 and 7%; the corresponding range for the percentages of supra-normal Z-scores were 0–13.2% and 0–9%, respectively, see Supplementary Figure H-A and H-B. There were no significant diagnostic differences in the percentage of female individuals with infra- or supra-normal regional values at baseline and follow-up ($P>0.05$). For infra-normal deviance the regions with highest percentages of change in female individuals with schizophrenia were the bilateral superior temporal and occipital regions with a decrease over time of 2.6% and an increase of 2.6%, respectively. For supra-normal deviance the bilateral superior temporal and occipital regions increased with 5.3% and decreased with 7.9%, respectively (see Figure H-C).

We assessed whether using z-scores from normative modeling led to stronger diagnostic group results as compared to using 'raw' morphometric similarity (i.e., the traditional approach). Therefore we averaged 'raw' morphometric similarity across the regions belonging to each of the functional networks. To investigate the diagnostic differences of 'raw' morphometric similarity scores for each functional network, we used seven separate generalized additive models (one for each functional network) to residualize the 'raw' morphometric similarity for age (included as a smooth term to allow for the potentially non-linear association between morphometric similarity and age), sex, Euler number, and scanner (Wood, SN, 2006). For the diagnostic comparison of the residualized 'raw' morphometric similarity we used Welch t-tests.

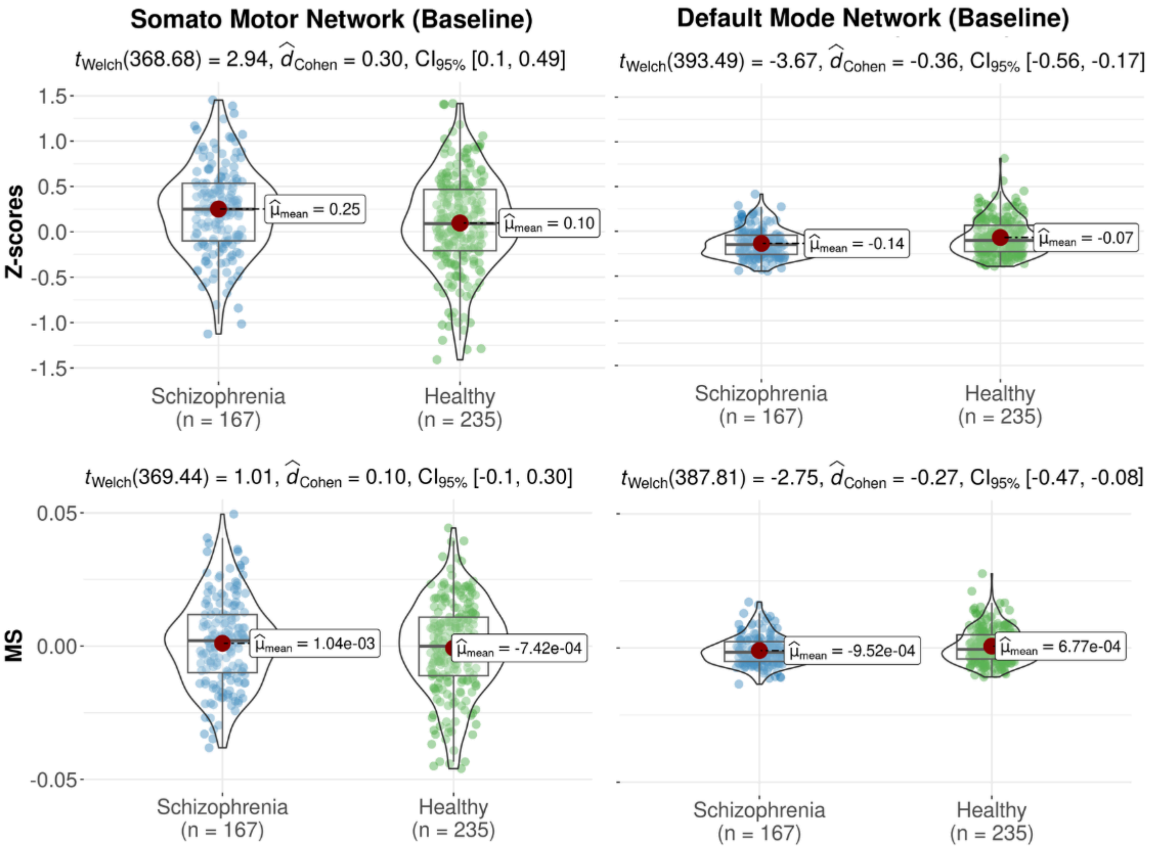


Supplemental Figure H. Violin plots of significant (FDR-corrected) differences in z-scores and ‘raw’ morphometric similarity (i.e. traditional values rather than normative modeling-based z-scores) of two functional networks between the group of healthy controls and the group of individuals with schizophrenia. MS, morphometric similarity.

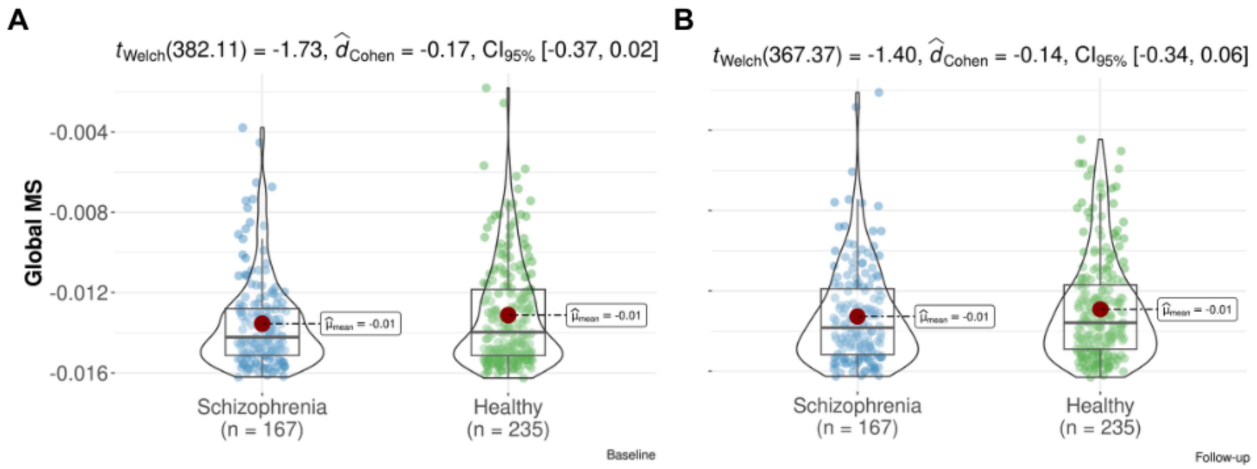


Supplemental Figure I. Violin plots showing differences in global morphometric similarity between the group of healthy participants and the group of individuals with schizophrenia for A) baseline, B) follow-up. Effect sizes are given either as Cohen's d. CI, 95% confidence interval. There were no statistically significant differences between diagnostic groups (baseline P=0.08, follow-up P=0.16).

Hall/CRC.
